# Phylo-HS: A phylogenetic hierarchical softmax for taxonomic classification

**DOI:** 10.1101/2025.01.27.634943

**Authors:** Romain Menegaux

## Abstract

Taxonomic binning –assigning taxonomic labels to DNA sequencing reads – is a core component of metagenomics data analysis. While machine learning approaches offer competitive speed and accuracy, their scalability is hindered by the growing number of referenced genomes and species. A major bottleneck lies in the final softmax layer of neural network models, which computes probabilities and gradients for all outcome classes. To address this, we propose **Phylo-HS**, a hierarchical softmax method that leverages the taxonomic tree to group classes into meaningful clusters. Phylo-HS achieves an order of magnitude speed improvement on a dataset with 5,000 classes and improves classification accuracy compared to frequency-based hierarchical softmax methods. By integrating phylogenetic structure into the model, Phylo-HS effectively balances scalability and accuracy for large-scale metagenomic analysis.

## 1 Introduction

With its drastic improvement in cost and efficiency, DNA sequencing has become a prime tool to study microbial communities. Modern DNA sequencing machines – so-called Next Generation Sequencing (NGS) – typically output up to the billions of short DNA sequences – *reads* – in random order. *Taxonomic binning* [Mande et al., 2012] is the task of assigning each of these reads to a taxonomic unit – a bacterial species for example. To do so reads are compared to an annotated reference genome database. As sequencing experiments are becoming commonplace, this reference database is becoming increasingly large, with new genomes assembled from sequencing experiments being uploaded every day [Szarvas et al., 2020]. Furthermore, our knowledge of the tree of life is constantly expanding with the discovery of new species [Parks et al., 2018], which complicates the task of taxonomic classifiers.

Standard approaches to taxonomic binning can be roughly divided into three categories. The first are alignment-based methods, that use string-matching algorithms to align reads to a selection of reference genomes with sequence alignment software such as BWA-MEM [Li, 2013] or Bowtie [Langmead et al., 2009]. The second category of methods, pseudo-alignment [Wood and Salzberg, 2014]; [Ounit et al., 2015]; [Kim et al., 2016]: [Bray et al., 2016], build databases of long discriminative *k*-mers ^1^ and store the taxa they are seen in. The *k*-mers from a read are looked up in this database and, if matches are found, the corresponding taxa are returned. Finally the third class of methods formulate taxonomic binning as a machine learning classification problem [Wang et al., 2007]; [McHardy et al., 2007]. In particular, large-scale linear methods [Vervier et al., 2016] and neural network classifiers [Menegaux and Vert, 2019]; [Liang et al., 2020] are competitive on modern datasets. Although these methods are fast, both their training and testing times are affected by having a large number of output classes.

The bottleneck with neural networks with large output spaces – well known in neural language modeling where the outcome classes are the whole word vocabulary – is the normalization of the class scores into probabilities, typically done with a softmax function. The computational complexity of this normalization step scales linearly with the number of output classes, and can make up a significant portion of training and testing times [Jozefowicz et al., 2016]. Several approaches have tackled this problem, either by (i) approximating the normalization factor by importance sampling [Bengio and Senécal, 2003]; [Mikolov et al., 2013b]; [Ji et al., 2016], (ii) replacing the softmax by self-normalized losses [Gutmann and Hyvärinen, 2010]; [Mnih and Teh, 2012] and (iii) replacing the softmax by an approximation such as the hierarchical softmax (HSM, [Goodman, 2001], [Morin and Bengio, 2005]).

The HSM groups outcome labels into clusters and decomposes classification in steps: first predicting the cluster and then choosing the label from that cluster. Introducing more levels in the hierarchy can improve the computational complexity up to the order 𝒪 of (*T/* log_2_(*T*)) compared to the regular softmax, where *T* is the number of labels or taxa. However the classification accuracy does slightly drop off, probably due to error propagation down the hierarchy levels. Designing the class hierarchy is primordial to mitigate this issue [Mnih and Hinton, 2009]. On the one hand, some approaches group classes according to their frequency, placing more common classes higher up in the tree, which has the double effect of speeding up computation and increasing the accuracy for these classes [Mikolov et al., 2011a]; [Mikolov et al., 2013a]; [Grave et al., 2017]. Another intuitive approach is to group the classes by similarity [Mnih and Hinton, 2009], so that the clusters themselves are informative. This has shown better classification accuracy than simply grouping them by frequency [Zweig and Makarychev, 2013]; [Chen et al., 2016].

Objectively defining class similarity can be challenging for some domains, however in taxonomic binning there is a natural hierarchy between classes: the phylogenetic tree. We take advantage of this structure in a new loss, Phylo-HS, which incorporates the phylogenetic tree in the hierarchical softmax. The contributions of this work are as follows:

- We give a simple algorithm to adapt the taxonomic tree to the hierarchical softmax.
- We show in experiments on large-scale datasets that the resulting loss outperforms the frequency-based HSM in classification accuracy, while maintaining the same or even slightly better classification and training speed.

After briefly reviewing related work in section 2, we will present the hierarchical softmax model and our method in detail in section 3. We present the experimental setup and results in section 4.

## 2 Related Work

We first review what initially motivated this work: the approximation of the softmax layer and the hierarchical softmax in particular. We then give an overview of other methods using a taxonomic tree for hierarchical classification.

### 2.1 Approximating the softmax

In neural network terminology, the softmax layer is a linear classifier: the composition of the softmax function with a linear operator *h → w · h*. Effectively it gives scores for each class *t* then normalizes them to sum to 1. In all that follows, we will write *h* as the *d*-dimensional input to the softmax layer, and *w* ∈ ℝ^*T ×d*^ as its parameters.

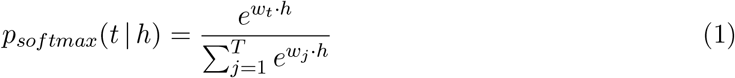

The denominator in (1) is also called the partition function *Z*(*h*; *w*). Computing this term has a complexity proportional to the number of output classes and the input dimension 𝒪 (*dT*). We now give the main lines of the many approaches that have been proposed to reduce this, most of which are reviewed in [Chen et al., 2016].

#### Reducing the effective input dimension

*d* Some methods have aimed at reducing the complexity of the individual dot products *w*_*j ·*_ *h*. The differentiated softmax [Chen et al., 2016] or the adaptive softmax [Grave et al., 2017] both assign a lower capacity *w* ∈ R^*d′*^, *d*^*′*^ *< d* to infrequent labels. The input *h* is then projected to a lower dimension before computing the dot products. If *T* ^*′*^ labels are assigned this smaller representation, the complexity of computing *Z* and its gradients is reduced to 𝒪 (*d*^*′*^*T* ^*′*^ + *d*(*T* − *T* ^*′*^)).

#### Approximating the partition function

Another popular approach is to approximate the denominator by importance sampling [Bengio and Senécal, 2003];[Mikolov et al., 2013b];[Ji et al., 2016]: the sum in *Z* is taken over a random subset of the samples instead. Although these methods do speed up training, at test time the true softmax must still be evaluated.

#### Self-normalized classifiers

[Devlin et al., 2014] maximize the unnormalized softmax likelihood with an added penalty for the normalization term (log *Z*)_2_. At test-time only the unnormalized scores are computed, forgoing the normalization completely. Other methods such as Noise Contrastive Estimation (NCE) [Gutmann and Hyvärinen, 2010]; [Mnih and Teh, 2012] change the loss function altogether and choose a binary classifier that distinguishes between the real training distribution and a noise distribution.

#### Hierarchical Softmax

The hierarchical softmax model (HSM) is an approximation to the softmax function that groups labels into clusters. When given an input *t*, the model predicts its cluster *c* with a first classifier *p*(*c* |*h*), then predicts the label *t* with a sub-model *p*(*t* |*c, h*). Since each label *t* belongs to a unique cluster *c*(*t*), the likelihood of *t* is the product

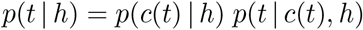

If the clusters are well balanced, this reduces the complexity of the softmax from 𝒪 (*dT*) to 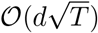. Introducing more levels in the hierarchy further lowers the complexity, up to 𝒪 (*d* log *T*) for a balanced binary tree [Morin and Bengio, 2005]. The assignment of labels to clusters is usually based either on their similarities [Mnih and Hinton, 2009] or on their frequencies [Mikolov et al., 2011b], with the former exhibiting higher accuracy [Chen et al., 2016]. We review the HSM in more detail in section 3.1.

Although much faster than the standard softmax, the HSM comes at a price in accuracy [Mikolov et al., 2011b]; [Zweig and Makarychev, 2013]. A solution proposed in [Le et al., 2011]; [Grave et al., 2017] strikes a balance between the hierarchical speedup and the softmax accuracy by introducing a short-list, treating frequent labels as single-label clusters.

### 2.2 Leveraging label taxonomy for classification

#### In machine learning

We focus here on classification problems in which the outcome classes are organized in a known tree hierarchy. This could in theory give extra information for classification which is not taken into account by traditional “flat” classifiers. In all generality the classes can be in the internal nodes in the tree, which calls for strategies to make predictions at different levels. [Silla and Freitas, 2011] give an extensive review of hierarchical classification across a variety of fields. More recently, [Wehrmann et al., 2018] designed a neural network architecture specially suited for multi-label classification. In the cases where the classes of interest are only in the leaf nodes of the tree, [Wu et al., 2017] give a hierarchical loss that penalizes errors differently depending on how far in the tree the prediction is from the true label. For instance predicting “dog” instead of “cat” is more harshly penalized than predicting “skyscraper”.

#### Taxonomic binning in metagenomics

In the metagenomics problem we are interested in, outcome classes are microbial taxa (such as bacterial species) which are organized in a taxonomic tree. Ideally predictions at finer levels such as species or genus are more desirable but making predictions at coarser levels such as phylum or family can also be of value. Indeed, some short DNA fragments can be shared by genomes of multiple species, in which case a prediction at a higher level can represent this ambiguity. Taxonomic binning software such as Kraken [Wood and Salzberg, 2014] and Centrifuge [Kim et al., 2016] both use the taxonomic tree to resolve multiple possible matches by returning the Least Common Ancestor (LCA) of all the matching taxa. Another way to use the taxonomic tree at test time is to sum estimated species-level probabilities to make predictions at the genus level [Busia et al., 2018]. This simple approach can be used by any binner outputting probabilities. In the most sophisticated approach yet, [Rojas-carulla et al., 2018] introduce a cascading softmax with separate softmax layers classifying each taxonomic rank *r*. Each softmax layer takes as input the hidden representation *h* and the output of the previous softmax layer. The loss function is a weighted average of the output softmaxes. This however does not reduce the computational complexity and on the contrary scales with the total number of nodes in the taxonomic tree rather than just the leaves.

## 3 Methods

### 3.1 Hierarchical softmax

We suppose established a tree hierarchy over *𝒯* labels. Reusing the notations in [Silla and Freitas, 2011], for a given vertex *t*∈ *𝒯*, we define ↑ (*t*) ∈ 𝒯as its parent, and ↓ (*t*) as the set of its children. We further denote *r*(*t*) ∈ ℕ as its depth – its distance to the root *t*_0_. With these notations, applying the function ↑ *r*(*t*) times to any vertex *t* returns to the root: ↑^*r*(*t*)^ (*t*) = *t*_0_.

For each internal node *t*, the hierarchical softmax model learns a classifier 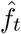 that distinguishes between its children given an input *h* ∈ ℝ^*d*^. Each of these classifiers is a softmax model with a |↓(*t*)| *× d* matrix of weights *w*:

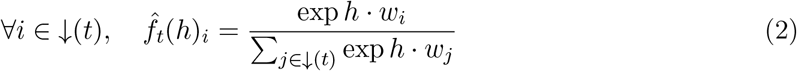

The computational complexity of each 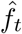therefore depends on its number of children 𝒪 (*d* | *↓* (*t*) |).

Under this model, the likelihood of any given node is the product of the conditional probabilities of all the nodes in its lineage:

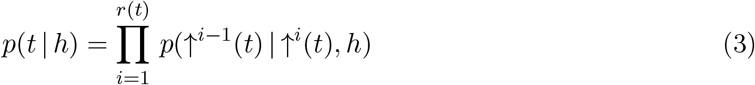

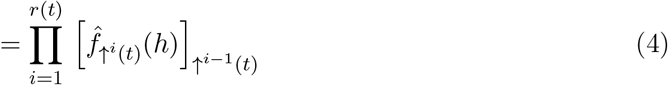

These parameters are learned jointly by maximizing the log-likelihood of the taxa in the training set (usually the leaves) through stochastic gradient descent (SGD).

#### Computational complexity

If the average number of children per ancestor node is *C*, the number of operations in the computation of the likelihood and gradient per sample *t* is 𝒪 (*d ×r*(*t*) *× C*). The quantity *r*(*t*) × *C* is minimized for binary trees, where it drops to log_2_(*T*) on average. One can further reduce the average *r*(*t*) by placing more frequent labels higher up in the tree. [Mikolov et al., 2013a] push this to the limit by Huffman coding the labels, which ensures an optimal average path length w.r.t. the empirical distribution. This tree, which we refer to as the Huffman tree, is also the one used in the hierarchical softmax loss of <monospace>fastText</monospace> [Joulin et al., 2016].

A detail perhaps worth mentioning is that binarizing the tree lowers the number of parameters for the HSM. In the non-binary version, a vector of *d*-dimensional weights is learned for each node except the root, so the total number of parameters for the HSM is (|𝒯 |− 1) × *d*. For the binary tree, this is reduced to (*T* − 1) ×*d* as each local classifier 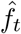 is a binary logistic model. Compared to the worst case scenario (a ternary tree), this corresponds to a reduction of 33%.

It is important to note that, at test-time, computing the probability for every class will still be an ℴ (*T*) operation. However if one is interested only in the top few most probable classes, some parts of the tree can be bypassed, as shown in figure 1. This is a factor to consider when designing the hierarchy 𝒯. For instance in the Huffman tree probabilities are computed for the most frequent labels first. Building a hierarchy based on similarity can also boost this by funneling the probability structure, rather than dispersing the probabilities higher up in the tree.

**Figure 1.**
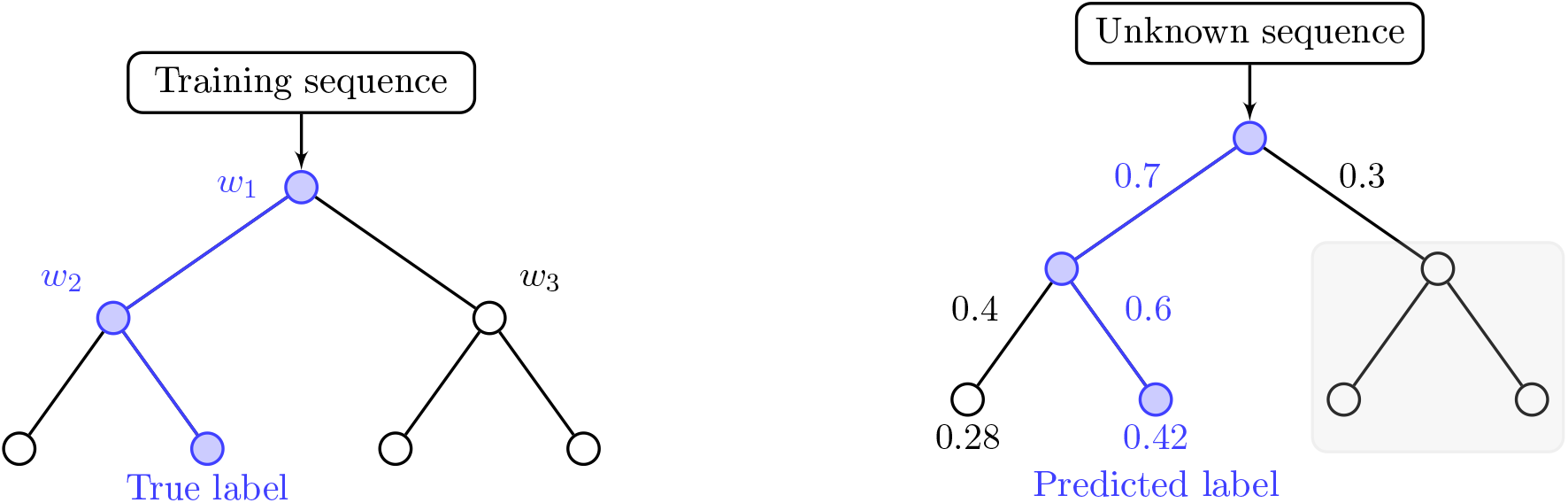
Example of a simple binary classification tree. On the left, we highlight the traversal of the tree for a training example with a known label. Only the weights *w*1 and *w*2 are used and updated during gradient descent. *w*3 is not used at all. On the right we show predicting the most probable label for a test sequence. Leaf probabilities are computed in a depth-first traversal. We show the state here after the probabilities on the left side are computed. Since the best label has a greater probability 0.42 *>* 0.3 than the right child of the root, the deeper children on the right do not have to be computed (shaded area)

### 3.2 Tree construction

#### Extracting from the reference tree of life

The classification tree for Phylo-HS is constructed at training time from the list of taxonomic IDs of training genomes, and a reference tree of life *𝒯* _*ref*_, ideally containing all species of interest. For each taxon *t* in the training set, we query the reference tree to get its full *lineage*, its path to the root *l*(*t*) = [*t, ↑(t)*, …, *↑*^*r(t)*^*(t)*] The new tree 𝒯 _*hs*_ is the minimal tree covering all training taxa and their ancestors. We further simplify *T*_*hs*_ by removing all nodes that have only one child.

#### Binarizing the taxonomic tre

In order to attain the maximal exponential speedup, we transform the resulting tree into a *full* binary tree, in which each node has either 0 or exactly 2 children. As shown in figure 2, leaf nodes are left untouched and nodes having only one child are removed. For nodes that have more than 2 children, we split the siblings into two separate groups according to their frequencies (total length of their representative genomes in training set). The *C* children are ordered by descending frequency [1, …, *C*] and then split into two groups [1, …, *K*] and [*K* + 1, …, *C*], where *K* is the first child such that |1| + … + |*K*| *>* |*K* + 1| + … |*C*|. This ensures that the total frequencies of each group are balanced, and that more frequent classes are on average higher up in the tree, hence accessed more rapidly. As the taxonomic hierarchy is conserved, the resulting tree can be seen as a hybrid between the Huffman coding tree and the taxonomic tree.

**Figure 2.**
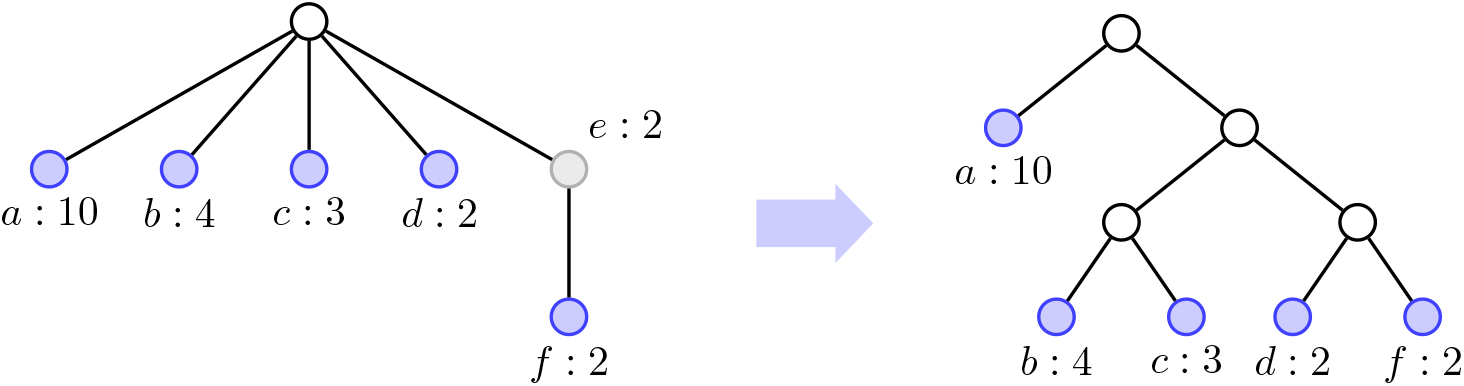
Example of binarization by frequency. The leaf taxons *a*; *b*; *c*; *d*; *f* are highlighted in blue, with the number of occurrences in the training dataset. For Phylo-HS, these frequencies are proportional to the total amount of bp belonging to genomes with taxon *a*; … ; *f*. On the *left*, they are laid out as originally. Taxon *e* has only one child and therefore is removed from the tree. On the *right*, the taxons as appearing in the final binary tree. Both children of each node should have a balanced total frequency.

## 4 Results

### 4.1 Learning the input representation

We test Phylo-HS on a taxonomic binning problem, where taxa must be assigned to short (∼50 − 400bp-long) DNA reads. A DNA read of length *L* is a sequence **x** = *x*_1_ … *x*_*L*_ ∈ 𝒜^*L*^, where 𝒜 = `{A, C, G, T}` is the alphabet of nucleotides. To transform a DNA read into an input representation *h* ∈ ℝ^*d*^, we use `fastDNA` [Menegaux and Vert, 2019]. More specifically, we segment a read into overlapping *k*-mers – contiguous subsequences of length *k*. A *d*-dimensional embedding is learned for each of the *N k*-mers, stored in an *N × d* embeddings matrix 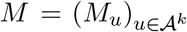. The representation *h* = Φ^*M*^ (**x**) of a read **x** is then simply the average of the embeddings of its constituent *k*-mers:

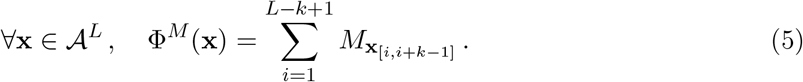

The embeddings matrix will be jointly trained by SGD with the output layer, either the full softmax layer or the HSM presented above. Training reads are extracted on the fly from genomes in the training set, with the addition of some random noise in the form of substitutions at a uniform rate.

### 4.2 Data

We test our method on 6 datasets of increasing complexity, composed by downloading genomes from the NCBI RefSeq database [NCBI Resource Coordinators, 2016]. The first 3 are *small, medium* and *large* from [Vervier et al., 2016], with respectively 51, 193 and 771 species represented in 356, 1564 and 2768 complete genomes. We will also use the validation set for *large* – composed of 193 genomes not present in the training set but whose species are – to evaluate the classification performance of the models. While the *large* dataset gave a comprehensive picture of the known bacterial and archeal genomes at the time of its publication in 2015, reference databases have since expanded.

As a more realistic dataset we use the 10,857 genomes of 3,640 species from the DeepMicrobes preprint [Liang et al., 2019]. It was obtained by filtering out similar species of the RefSeq database, by comparing the similarity of one of their representative genomes. Our largest dataset is *RefSeq*

*CG* from the benchmark [Ye et al., 2019]. It contains all the complete microbial genomes available in RefSeq as of 2018, a total of 23000 genomes belonging to 13000 taxa. As a large portion of those taxa are viruses having relatively small genomes, we simplify this dataset by attributing the same domain-level taxonomic id (label) to all viruses. The resulting dataset contains the same number of genomes with 5500 taxa.

As a reference taxonomic tree, we use the NCBI taxonomy downloaded on 30/09/2020.

### 4.3 Classification accuracy

The measures reported here are of species-level recall and precision. For a given species *t*, if *n*_*true*_ is the number of validation sequences of species *t, n*_*pred*_ is the number of sequences classified as *t* by the model, and finally *n*_*tp*_ is the number of true positives: sequences correctly classified as *t*. Recall (or sensitivity) for a species is the proportion *n*_*tp*_*/n*_*true*_. Average species-level recall is obtained by averaging this value over all the species truly present in the validation set. Precision is the proportion *n*_*tp*_*/n*_*pred*_. Average species-level precision is obtained by averaging this value over all the predicted species.

In figure 3, `fastDNA` models were trained for the same number of epochs on the *large* dataset then evaluated on the *large* validation set. One epoch corresponds to a number of reads sufficient to cover on average once all the positions in the training genomes. Phylo-HS does better both in terms of precision and recall than the Huffman coding alternative, although the gap does narrow as the model capacity gets higher (larger *k*). Similar results have been reported for language models [Mnih and Hinton, 2009]; [Chen et al., 2016]. This does not however fully compensate the performance dropoff of the hierarchical softmax compared to the full softmax.

**Figure 3.**
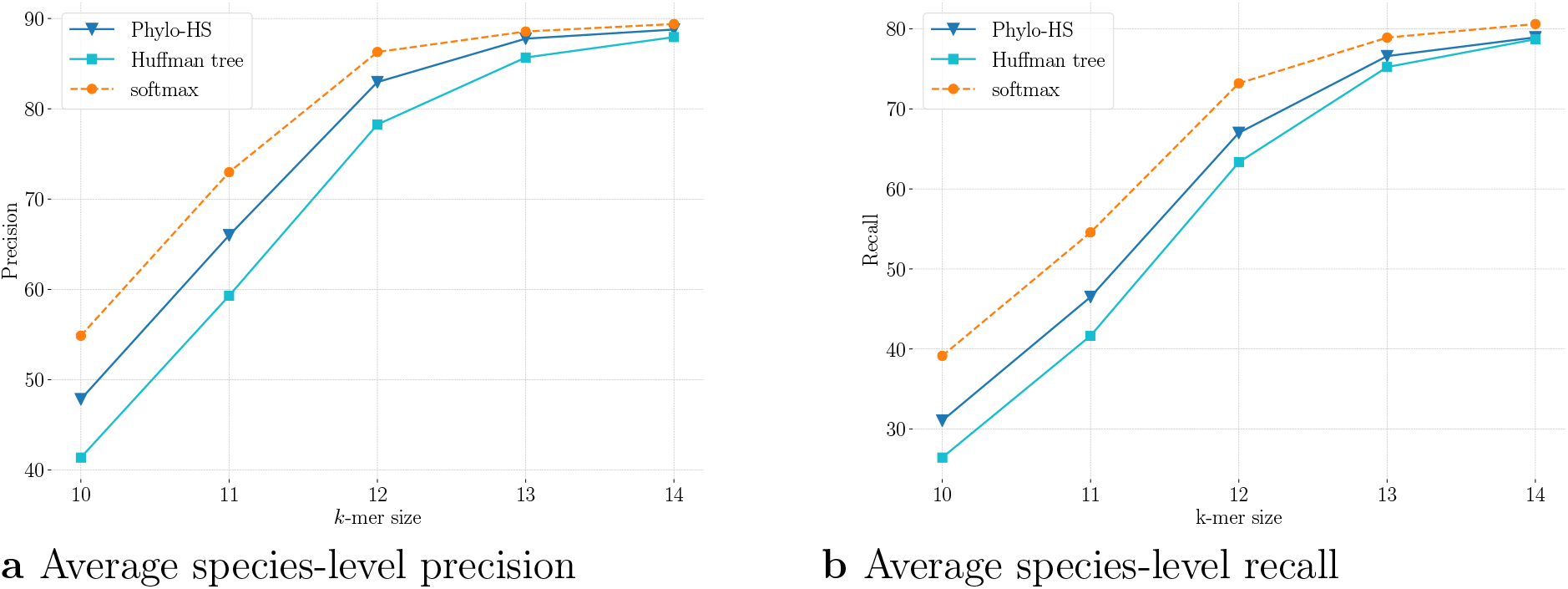
Comparison of the effect of the loss function on the classification performance of a `fastDNA` model, on the *large* dataset. Results are shown as a function of the *k*-mer size *k*. The models, all of embedding dimension *d* = 50, were trained for 100 epochs with a learning rate of 0.1 and a mutation noise rate of 4%.

### 4.4 Classification speed

All speed tests were performed on a Intel Xeon CPU E5-2440 0 - 2.40GHz with 12 cores and 128GB of RAM. Models were trained using 12 threads, and tested with a single thread. There was some variability in our speed measurements, so we report the median time over 10 runs. Models all have the same hyperparameters *k* = 14 and *d* = 50 and were trained for 100 epochs. We do not include fixed costs such as model loading time to concentrate solely the asymptotic per-read speed. These fixed costs are the same for the studied models.

As confirmed in figure 4, training the full softmax for the larger label sets becomes prohibitively slow, while the hierarchical softmax models are much less affected. Both versions of the hierarchical softmax are equally fast in both training and testing. While there is perhaps a slight edge for Phylo-HS at testing time, this could be attributed to variance in the speed measurements or data-specific quirks. As expected, the speed improvement with respect to the full softmax does diminish between test and training, but remains a substantial 10*×* speedup on the largest dataset.

**Figure 4.**
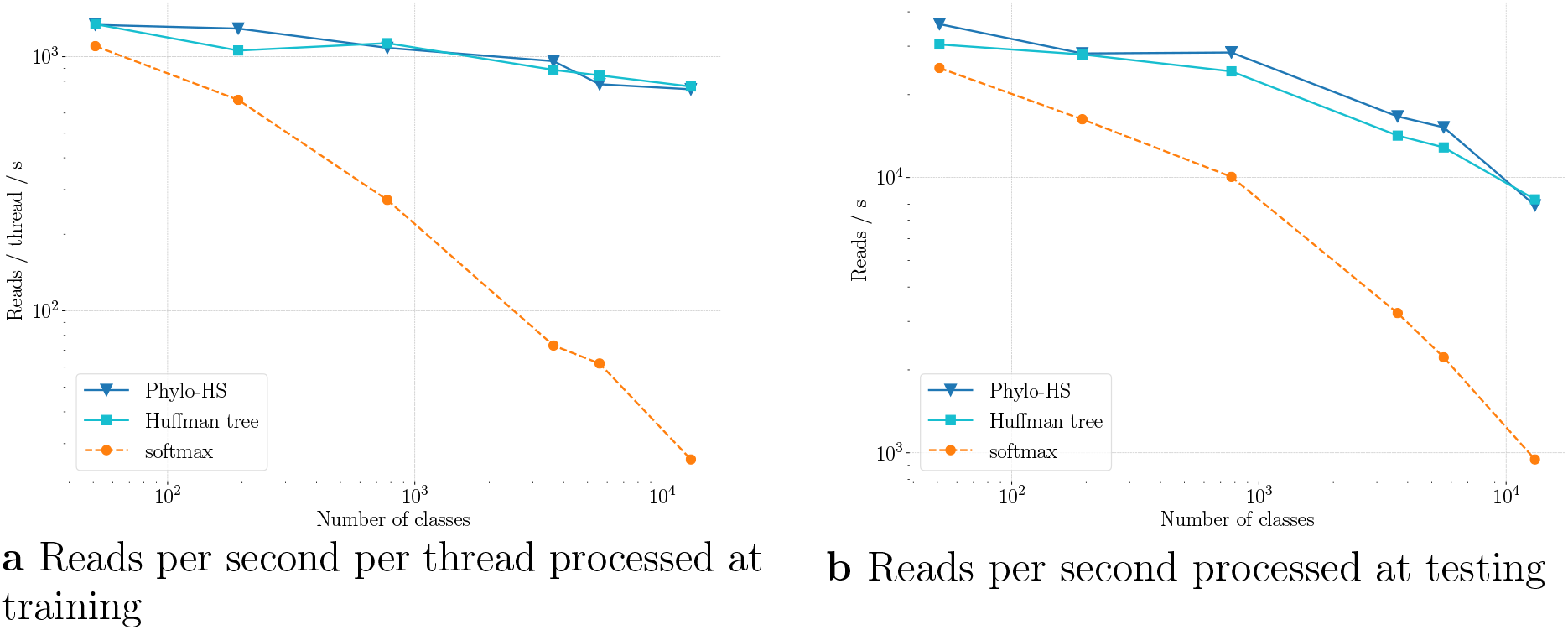
Comparison of the effect of the loss function on the number of reads per second processed by a `fastDNA` model. Results are shown as a function of the number of output classes. All models have *k*-mer size *k* = 14 and embedding dimension *d* = 50. Input reads are 200-bp long.

## 5 Discussion and future work

We have shown with these preliminary experiments that modeling the hierarchical softmax on the taxonomic tree can bring gains in classification accuracy at no cost in computational complexity. However it must be noted that the frequency-based trees such as the Huffman coding one do still have the advantage of being nomenclature-agnostic. Indeed they do not depend on labels being recognized as official taxonomic IDs, which makes them potentially better suited for characterizing genomes of unknown species. One could still apply Phylo-HS in that case by clustering genomes based on simple measures of similarity such as Average Nucleotide Identity (ANI) for instance. In fact many tools have been developed to infer phylogenetic trees from raw genomes [Ankenbrand and Keller, 2016]; [Na et al., 2018]; [Szarvas et al., 2020], and in particular for metagenome assembled genomes (MAGs) [Wu, 2018]. It could be a promising approach to use these tools on the training genomes to build the taxonomic classification tree.

## 6 Acknowledgements

I thank Jean-Philippe Vert for helpful discussions and comments.

